# The vaginal microbiota of pregnant women varies with gestational age, maternal age, and parity

**DOI:** 10.1101/2023.02.26.530121

**Authors:** Roberto Romero, Kevin R. Theis, Nardhy Gomez-Lopez, Andrew D. Winters, Jonathan Panzer, Huang Lin, Jose Galaz, Jonathan M. Greenberg, Zachary Shaffer, David J. Kracht, Tinnakorn Chaiworapongsa, Eunjung Jung, Francesca Gotsch, Jacques Ravel, Shyamal D. Peddada, Adi L. Tarca

## Abstract

The composition of the vaginal microbiota is heavily influenced by pregnancy and may factor into pregnancy complications, including spontaneous preterm birth. However, results among studies have been inconsistent, due in part to variation in sample sizes and ethnicity. Thus an association between the vaginal microbiota and preterm labor continues to be debated. Yet, before assessing associations between the composition of the vaginal microbiota and preterm labor, a robust and in-depth characterization of the vaginal microbiota throughout pregnancy in the specific study population under investigation is required. Herein, we report a large longitudinal study (N = 474 women, 1862 vaginal samples) of a primarily African-American cohort– which experiences a relatively high rate of pregnancy complications – evaluating associations between individual identity, gestational age, and other maternal characteristics with the composition of the vaginal microbiota throughout gestation resulting in term delivery. The primary factors influencing the composition of the vaginal microbiota in pregnancy are individual identity and gestational age at sampling. Secondary factors are maternal age, parity, obesity, and self-reported *Cannabis* use. The principal pattern across gestation is for the vaginal microbiota to remain or transition to a state of *Lactobacillus* dominance. This pattern can be mitigated by maternal parity and obesity. Regardless, network analyses reveal dynamic associations among specific bacterial taxa within the vaginal ecosystem, which shift throughout the course of pregnancy. This study provides a robust foundational understanding of the vaginal microbiota in pregnancy among African-Americans, in particular, and sets the stage for further investigation of this microbiota in obstetrical disease.

**IMPORTANCE:** There is debate regarding links between the vaginal microbiota and pregnancy complications, especially spontaneous preterm birth. Inconsistencies in results among studies are likely due to differences in sample sizes and cohort ethnicity. Ethnicity is a complicating factor because, although all bacterial taxa commonly inhabiting the vagina are present among all ethnicities, the frequencies of these taxa vary among ethnicities. Therefore, an in-depth characterization of the vaginal microbiota throughout pregnancy in the specific study population under investigation is required prior to evaluating associations between the vaginal microbiota and obstetrical disease. This initial investigation is a large longitudinal study of the vaginal microbiota throughout gestation resulting in a term delivery in a primarily African-American cohort, a population that experiences disproportionally negative maternal-fetal health outcomes. It establishes the magnitude of associations between maternal characteristics, such as age, parity, BMI, and self-reported *Cannabis* use, on the vaginal microbiota in pregnancy.

## INTRODUCTION

The composition of the vaginal microbiota is broadly consistent across populations of reproductive age women (1–5). In general, the vaginal microbiota can be categorized into five primary community state types (CSTs) that are defined by a predominance, or a lack thereof, of *Lactobacillus* spp. (1–9). Four of these CSTs are dominated by *Lactobacillus* spp. (*L*. *crispatus -* CST I, *L*. *gasseri* - CST II, *L*. *iners* - CST III, *L*. *jensenii* - CST V) and the other CST (CST IV) is typically not dominated by any one bacterium, but rather is comprised of a diverse array of microorganisms (4-6, 8-10). CST IV has been further subcategorized as CST IV-A or CST IV-B (11). CST IV-A is characterized by high relative abundances of *Candidatus* Lachnocurva vaginae (formerly Bacterial Vaginosis-Associated Bacterium 1, or BVAB1 (12)) *Gardnerella vaginalis,* and *L. iners*, whereas CST IV-B has high relative abundances of *Atopobium vaginae*, *G. vaginalis*, and *L. iners* (5, 11). Importantly, the *Lactobacillus*-dominated CSTs (I, II, III, V), and especially CST I, which is dominated by *L. crispatus*, are associated with optimal vaginal health (13–19) and positive reproductive outcomes (20–29). In contrast, CST IV-A and CST IV-B have been associated with bacterial vaginosis (7, 13, 30–35) and, among pregnant women, CST IV (23, 25, 36–38), CST IV-associated bacteria (23, 25–29, 38–41), and/or a greater vaginal microbiota diversity in general (22), have been associated with an increase in the risk of spontaneous preterm birth (sPTB) – the leading cause of neonatal mortality and mobility worldwide (42, 43). Nevertheless, non-pregnant and pregnant women alike with vaginal microbiotas classified as CST IV can be asymptomatic (4, 44), and their reproductive health and pregnancy outcomes are generally normal. Therefore, the strength and clinical relevance of associations between vaginal CSTs and female reproductive health and pregnancy outcomes remains unclear (45).

Ethnicity is a complicating factor in such studies as it is associated with the structure of the vaginal microbiota – all CSTs are present among all ethnicities, yet the frequencies of the CSTs among ethnicities vary (2-4, 22, 46-48). For example, African American and Hispanic women are more likely to exhibit CST IV vaginal communities, whereas Caucasian and Asian women tend to more frequently display *Lactobacillus*-dominated CSTs (2–4). Overall, regardless ethnicity, the composition of the vaginal microbiota can be highly labile and some of the factors influencing this lability include the menstrual cycle (5, 49–54), sexual activity (55–57), and pregnancy (58). The menstrual cycle appears to have a stabilizing effect on the composition of the vaginal microbiota, an affect that has been attributed to high estrogen levels, which favor the proliferation of *Lactobacillus* spp. (5, 49–54). Conversely, sexual activity increases the likelihood of CST IV vaginal communities (57) and decreases the presence of potentially protective *L. crispatus* (56). Pregnancy, a vulnerable period accommodating the growth and development of the fetus, and that includes a drastic rise in steroid hormones (e.g. progesterone and estrogen) (59, 60), also favors the presence of *Lactobacillus*-dominated CSTs in the vagina (44, 58). Indeed, we previously reported that the vaginal microbiota in pregnancy differs from that in non-pregnant women (58). Specifically, pregnant women have higher relative abundances of *L. vaginalis*, *L. crispatus*, *L. gasseri* and *L. jensenii*, and lower abundances of 22 other non-*Lactobacillus* phylotypes (58). In addition, the vaginal microbiota of pregnant women is typically more stable (i.e., consistent across time) than that of non-pregnant women (58). These general findings have been replicated by other investigators (20, 21, 48). Therefore, it has been proposed that increased stability of the vaginal microbiota and *Lactobacillus*-dominance during pregnancy play a protective role and reduce the likelihood of pregnancy complications, especially sPTB (47, 61). However, the association between variation in the composition of the vaginal microbiota and preterm birth continues to be debated (45, 62–64). Potential explanations for the inconsistencies in results among published studies include differences in sample sizes and cohort ethnicity. Therefore, before evaluating associations between the composition of the vaginal microbiota and obstetrical disease, including sPTB, a robust and in-depth characterization of the vaginal microbiota throughout pregnancy in the specific study population under investigation is required.

This initial investigation focuses on an urban population that experiences a high risk of pregnancy complications (65–75). It uses 16S rRNA gene amplicon sequencing to assess the trajectory of the composition of the vaginal microbiota throughout gestation ending in term delivery. Leveraging longitudinal samples from a large set of patients with well-characterized demographic and clinical data, this study establishes the magnitude of associations between maternal characteristics, such as age, parity, and ethnicity on the vaginal microbiota. Such knowledge is important for the assessment of previous reports and for informing future analyses of the vaginal microbiota in relationship to obstetrical complications, especially sPTB. Moreover, this study provides information on a primarily African American population, for which available data are overall sparse despite this population experiencing disproportionally negative maternal-fetal health outcomes (65–75).

## RESULTS AND DISCUSSION

The demographic characteristics of the 474 patients with term delivery included in this study (the largest cohort sampled to date) are presented in **Table 1**. This cohort is primarily African-American [94.5% (448/474)] with a body mass index (BMI) above 25 kg/m^2^ [65% (306/472)]. The distribution of the gestational ages at which the 1862 vaginal fluids were collected from these patients is depicted in **Figure 1**. Each woman had 3 to 4 samples (median of 4) collected between 8 and 38^+6^ weeks of gestation.

**Table 1.** Clinical and demographic characteristics of the study population. Data are given as median (interquartile range, IQR) and percentage (n/N). ^a^Two missing data

|  | **Term Birth**<br>**(n = 474)** |
| --- | --- |
| Maternal age (years; median [IQR]) | 24 (21-27) |
| Body mass index (kg/m<sup>2</sup>; median [IQR]) | 27.5 (22.7-33.7)<sup>a</sup> |
| Primiparity | 20.9% (99/474) |
| Race/Ethnicity |  |
| African-American | 94.5% (448/474) |
| White | 1.7% (8/474) |
| Asian | 0.2% (1/474) |
| Hispanic | 0.2% (1/474) |
| Other | 3.4% (16/474) |
| Gestational age at delivery (weeks; median [IQR]) | 39.6 (39-40.4) |
| Cesarean Section | 25.9% (123/474) |
| Fetal sex |  |
| Female | 48.7% (231/474) |
| Male | 51.3% (243/474) |
| Birthweight (grams; median [IQR]) | 3286 (3091-3580) |

**Figure 1.**
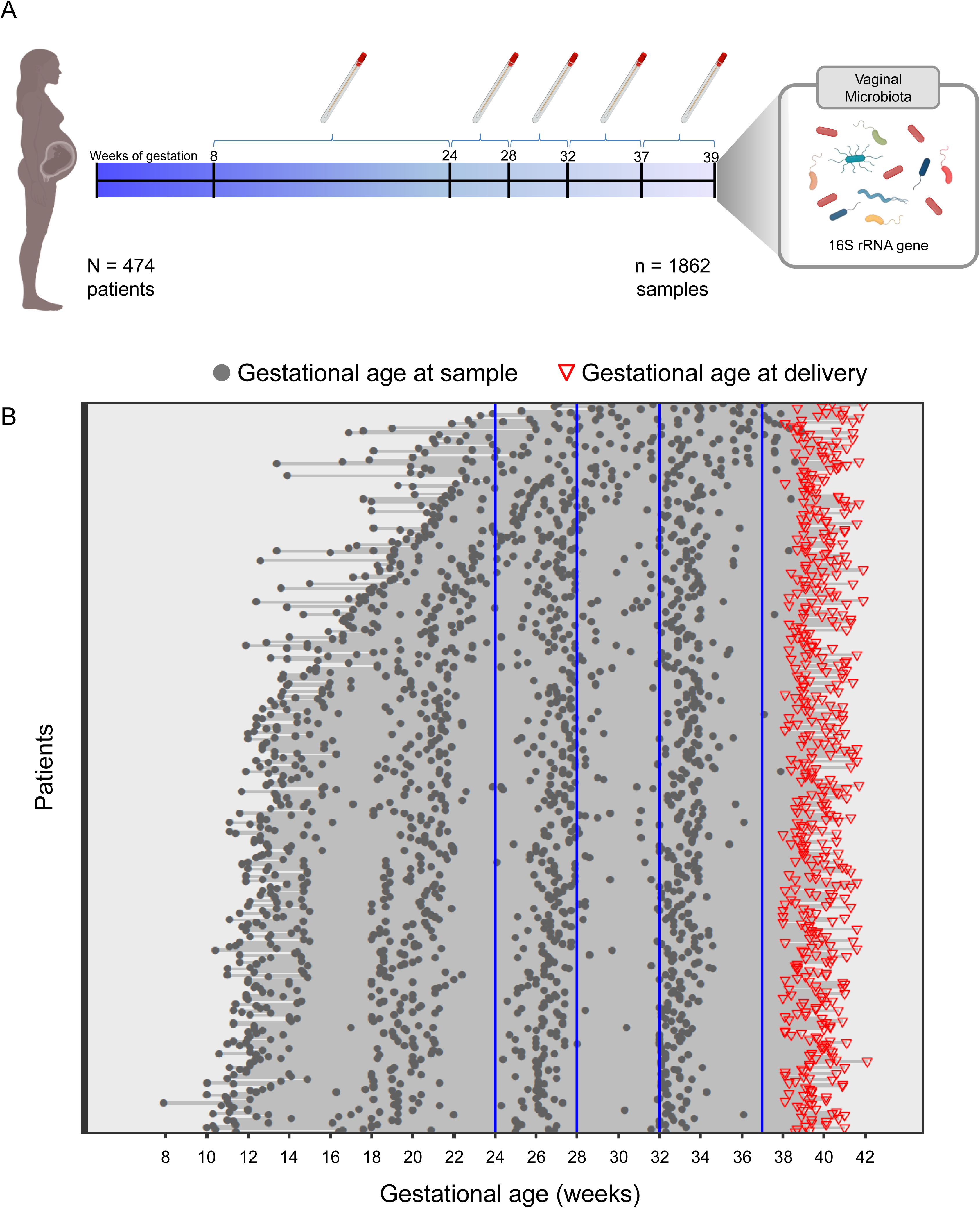
Gestational ages at the time of vaginal fluid sample collection in a cohort of women ultimately delivering at term. (A) 1,862 vaginal fluid samples were collected from 474 pregnant women between 8 and 38^+6^ weeks of gestation. The vaginal microbiota was profiled using 16S rRNA gene sequencing. (B) Each line corresponds to one patient and each dot to a sample for which the vaginal microbiota was characterized. Gestational ages at delivery are shown using red triangles.

### Effect of patient/subject identity

The structure of the vaginal microbiota during pregnancy has been reported to vary with gestational and maternal age (47, 48, 58). However, the scope and strength of all factors potentially influencing the structure and fluidity of the vaginal microbiota during pregnancy remain to be elucidated. In the current study, beta diversity, or the shared diversity of the microbiota between samples, was characterized using the Jaccard (i.e., microbiota composition) and Bray-Curtis (i.e., microbiota structure) indices. The variation in vaginal microbiota composition and structure was primarily explained by patient identity (SubjectID: composition - R^2^=58%-61%; structure - R^2^=65%-68%) and by the patient-specific variation with gestational age (interaction between SubjectID and Gestational age: composition - R^2^=16%-18%; structure - R^2^=14%-16%). Only relatively modest percentages of the variance in the composition and structure of the vaginal microbiota were explained by maternal characteristics, such as age (0.2%-1.4%), parity (0.3%-1.9%), and self-reported *Cannabis* use (0.3%-1.8%). Overall, these findings illustrate the large influence that patient (i.e., individual) identity has on the composition and structure of the vaginal microbiota, which is consistent with general observations of microbiotas at other body sites, such as the oral cavity, gut, and skin (76–81). Furthermore, this highlights the importance of robust longitudinal, as opposed to cross-sectional, studies to account for inter- and intra-individual variability in the microbiota (80, 82–84).

### Effect of gestational age

Alpha diversity, which is the diversity of the microbiota within individual samples, was characterized using Chao1 (i.e., richness) and Shannon and Simpson (i.e., evenness) indices. Both richness (**Figure 2A,C**) and evenness (**Figure 2B,D**) of the vaginal microbiota decreased with advancing gestational age from the first to the third trimester (p<0.0001 for all). This is consistent with previous reports of alpha diversity in cohorts of primarily African-Americans (28, 48, 58). In contrast, previous reports of largely Caucasian cohorts indicated that alpha diversity is generally low and consistent throughout the entirety of gestation (23, 47, 48, 85). Nevertheless, in the current study, there was substantial heterogeneity in the rate of decrease in vaginal microbiota alpha diversity among patients – the decrease was steeper for women who had higher baseline diversity early in pregnancy (correlation between random intercepts and random slopes: Shannon -0.79, Simpson -0.67, Chao1= -0.67) (see for example **Figure 3**). The decrease in alpha diversity with advancing gestational age remained significant after adjusting for potential confounding variables, including maternal age, parity, and BMI (**Table 2**). These alterations in the overall structure of the vaginal microbiota likely reflect physiological alterations (e.g., glycogen levels (5, 86, 87)) in the vaginal microenvironment across gestation that favor the predominance of a few bacterial taxa that can thrive in these conditions (e.g. *Lactobacillus* spp.).

**Figure 2.**
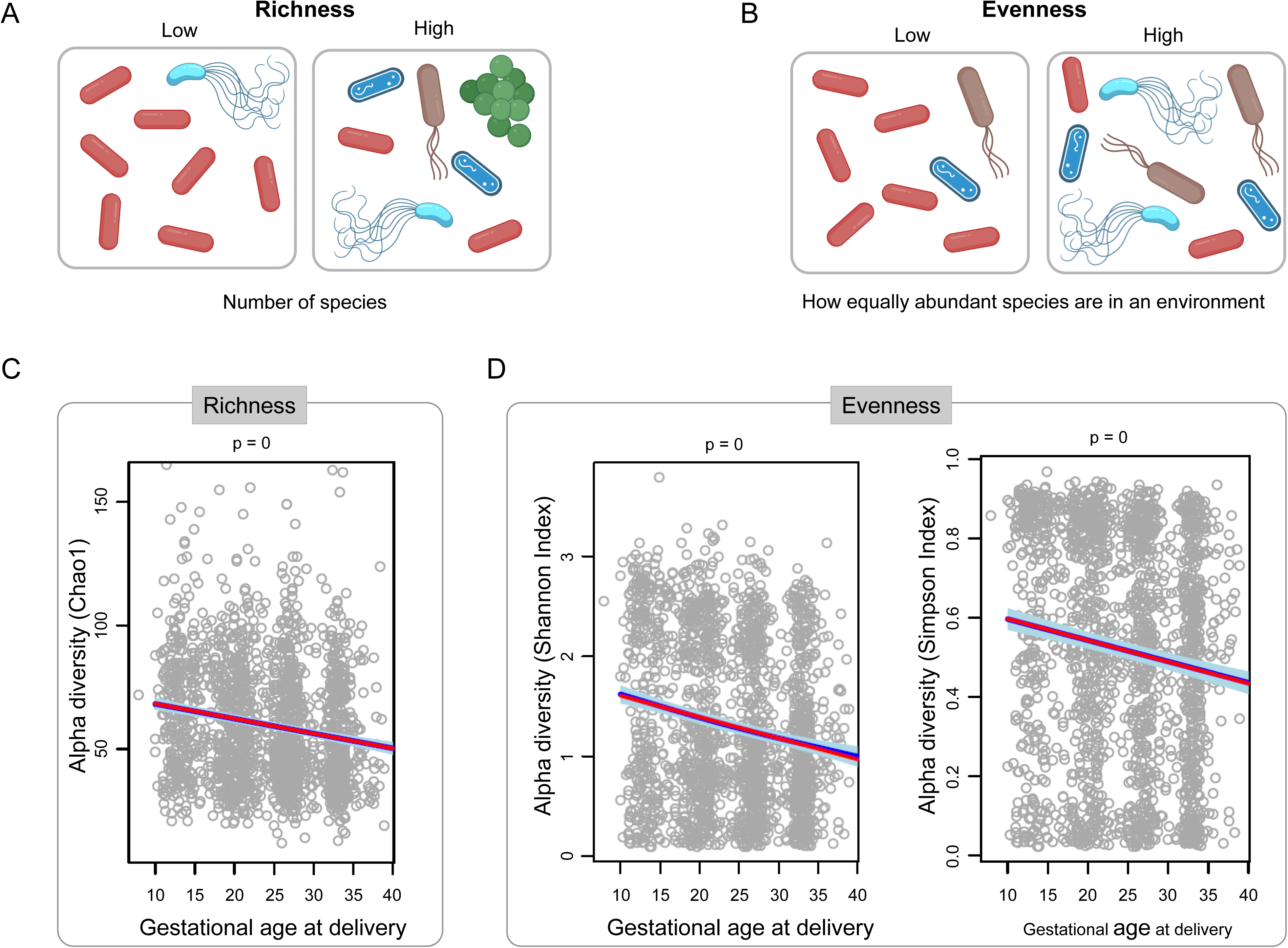
Decrease in alpha diversity of the vaginal microbiota with gestational age in women ultimately delivering at term. Graphical representation of low and high bacterial community richness (A) and evenness (B). Linear mixed-effects models illustrating decreases of bacterial community richness (C) and evenness (D) over the course of gestation. Each dot corresponds to one sample. The red line represents the linear fit using linear mixed-effects models. The dark blue line represents the model fit and light blue areas define the 95% confidence intervals derived from generalized additive models with splines transformation of gestational age.

**Figure 3.**
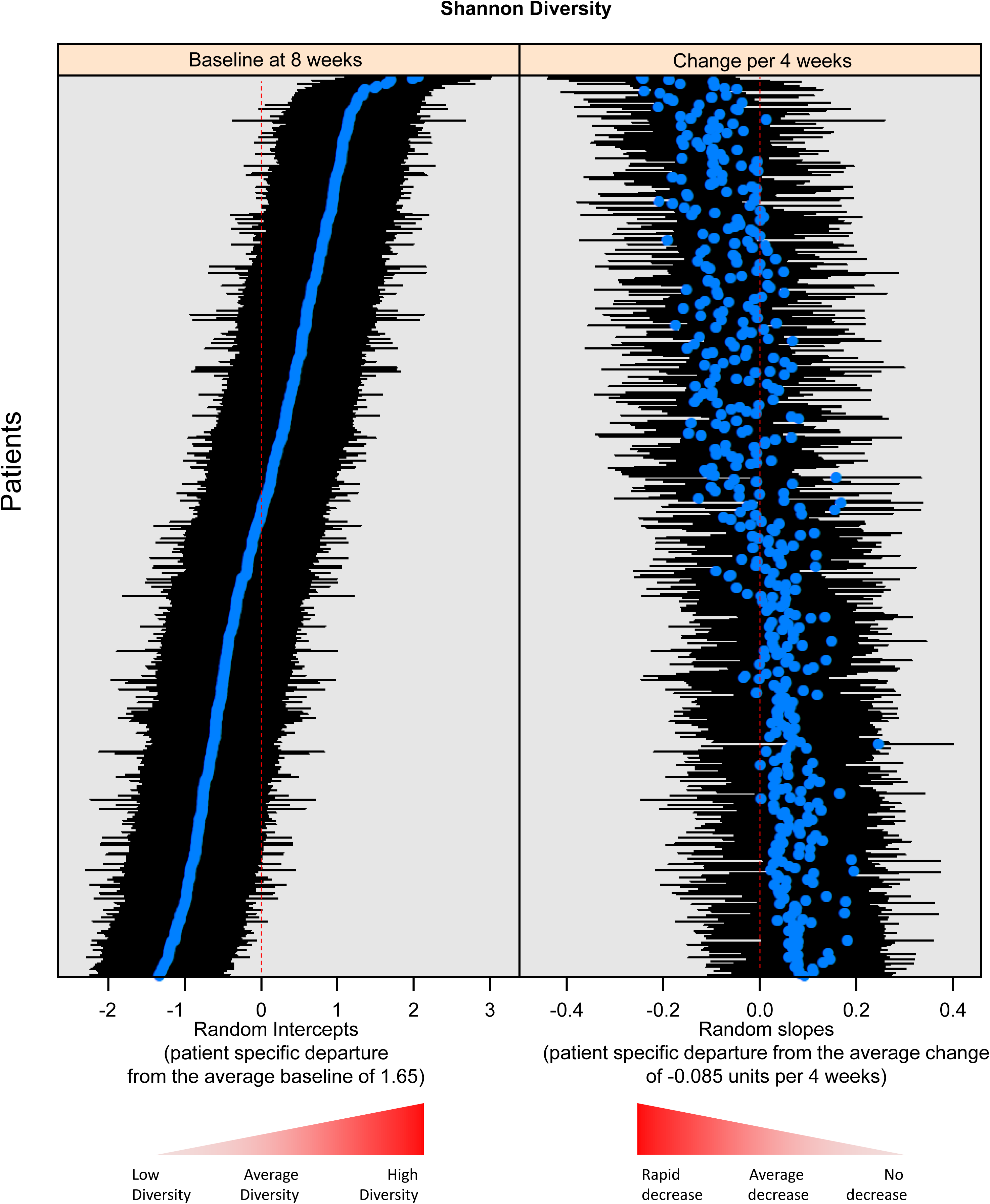
Rate of decrease in alpha diversity (Shannon diversity index) of the vaginal microbiota with gestational age is steeper in women with higher baseline diversity. The left panel shows the baseline diversity for each patient (blue dots) and corresponding 95% confidence intervals (black lines). The right panel shows the rate of change in diversity (blue dots) and confidence intervals (black lines). Women who had higher baseline diversity had steeper decrease in diversity with advancing gestation (correlation between random intercepts and random slopes of -0.79).

**Table 2.** Differences in alpha diversity values (Chao1 richness, Shannon, and Simpson diversity) of the vaginal microbiota profiles of women who delivered at term. *Gestational age was centered at 8 weeks and then scaled by 4, therefore the change in diversity corresponds to a four week interval **Maternal age was scaled by 5, therefore the diversity corresponds to a 5 year increase in maternal age

| Diversity | Covariate | Estimate | p |
| --- | --- | --- | --- |
| Chao1 | Gestational age* | -2.41 | 0.000 |
| Chao1 | Obese | 4.19 | 0.033 |
| Shannon | Gestational age* | -0.08 | 0.000 |
| Shannon | Parity | 0.09 | 0.000 |
| Shannon | Ethnicity Hispanic/Latino | -0.40 | 0.024 |
| Simpson | Gestational age* | 0.56 | 0.000 |
| Simpson | Age** | -0.02 | 0.000 |
| Simpson | Parity | 0.03 | 0.000 |
| Simpson | Ethnicity Hispanic/Latino | -0.23 | 0.003 |
| Simpson | Race other than African-American | 0.14 | 0.014 |

The vaginal microbiota is consistently categorized into CSTs that are defined by a dominance or lack thereof of *Lactobacillus* spp. (4, 11, 23, 44). Using a previously established protocol for assigning CSTs to vaginal samples based on 16S rRNA gene sequence data (11), we identified seven CSTs among the 1862 samples included in this study (**Figure 4A**). These CSTs included four dominated by *L. crispatus* (I), *L. gasseri* (II), *L. iners* (III), or *L. jensenii* (V), and three more diverse CSTs (CST IV) comprised of *L. iners*, *Gardnerella* sp., and *Megasphaera* sp., with Ca. Lachnocurva vaginae, *Atopobium vaginae*, and *Bifidobacterium* sp. being relatively abundant in CST IV-A, -B, and -C, respectively. These CSTs are consistent with previous investigations of the vaginal microbiota in smaller cohorts (27, 29, 36, 39–41, 44, 47, 58, 85, 88–94), further illustrating the depth of complexity of CST IV-designated communities.

**Figure 4.**
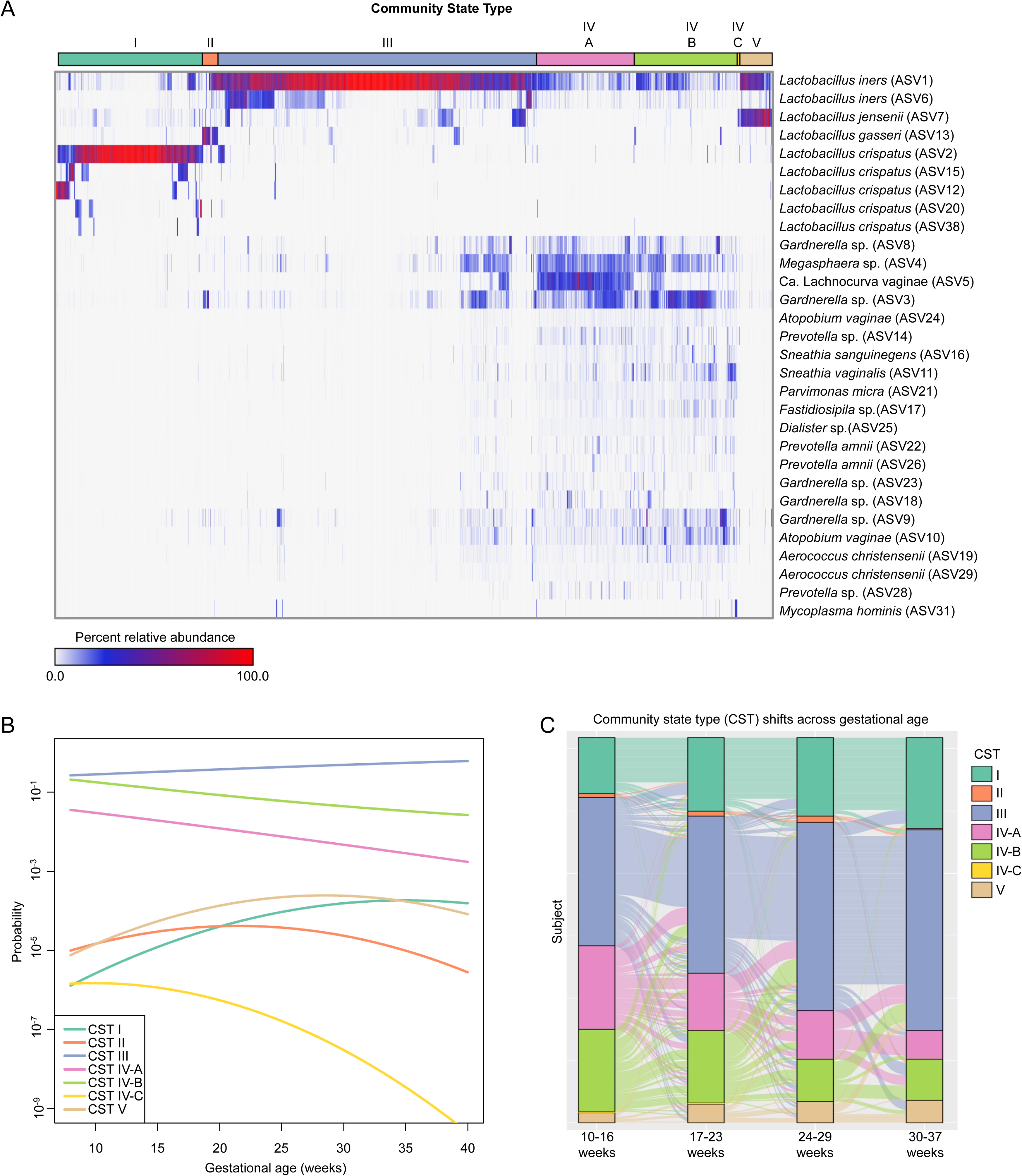
Variation in the community state type (CST) of the vaginal microbiota throughout gestation among women who ultimately delivered at term. (A) Heatmap illustrating the relative abundances of the 30 most abundant amplicon sequence variants (ASVs) among the vaginal 16S rRNA gene profiles. The bar on top indicates vaginal CSTs assigned using the program VALENCIA (11). (B) Dynamics of vaginal CST prevalence as a function of gestational age among women ultimately delivering at term. The log-odds of membership for each CST were modeled using binomial linear-mixed effects models. Fixed effects in these models included gestational age (linear and quadratic terms, as needed) and maternal characteristics, while one random intercept was allowed for each subject. (C) Alluvial plot illustrating the temporal dynamics of vaginal CST prevalence and transitions among 309 women who delivered at term and contributed one sample per each of the four discrete time points (10 to 37 weeks).

In this study, CST prevalence was a function of gestational age (**Figure 4B**). Except for the two least abundant CSTs (II and IV-C), for which statistical power was inherently limited, the membership probability to any CST displayed dynamic changes with gestational age (p<0.05 for all; **Figure 4B**). While *Lactobacillus*-dominated CSTs I, III, and V tended to be more abundant with advancing gestational age, the abundance of the more diverse CSTs IV-A and -B declined steadily as term gestation approached (**Figure 4B**). Notably, in a secondary analysis of women (N = 309) for which samples were available from each of four discrete time points across gestation, there was a pronounced shift in CST composition with advancing gestational age; specifically, there was an increase in CSTs I and III at the end of pregnancy, derived primarily from patients with an initial CST IV-A or IV-B (**Figure 4C**). These findings are in line with prior cross-sectional (21, 24, 29, 36, 39, 40, 48, 88, 89, 91, 93) and longitudinal (22, 23, 25–28, 41, 44, 47, 48, 58, 85, 90, 92, 94, 95) studies which included characterization of the structure and dynamics of the vaginal microbiota in term pregnancies. At a community level, pregnancy has been shown to create favorable conditions for a *Lactobacillus*-dominated vaginal microbiota, particularly CSTs I and III, and a shift away from the more diverse CSTs IV-A and IV-B, as gestation progresses to term (20, 23, 48, 58). Although shifts in the vaginal microbiota occur in both gravid and non-gravid women, an increased prevalence of specifically *Lactobacillus*-dominated CSTs in pregnant women is intriguing because it could protect against ascending infection (96), which could culminate in sPTB, through the competitive exclusion of opportunistic pathogens within the vaginal microenvironment (47, 97). Furthermore, *Lactobacillus* species produce lactic acid, which has anti-inflammatory properties (98–100). Additional research exploring the functional role of the vaginal microbiota on the host in large longitudinal cohorts is warranted to address these hypotheses.

After describing the changes in the composite measures of the vaginal microbiota alpha and beta diversity and CSTs, we utilized linear mixed-effects (LME) modeling to analyze the relationships between gestational age and maternal characteristics with the relative abundances of individual bacterial taxa denoted as amplicon sequence variants, or ASVs **(Supplemental Table 1)**. Increased gestational age was positively correlated with 33 exclusively *Lactobacillus* ASVs (q<0.1), and negatively correlated with ASVs that are typical members of vaginal CST IV **(Supplemental Table 1, Figure 5)**.

**Figure 5.**
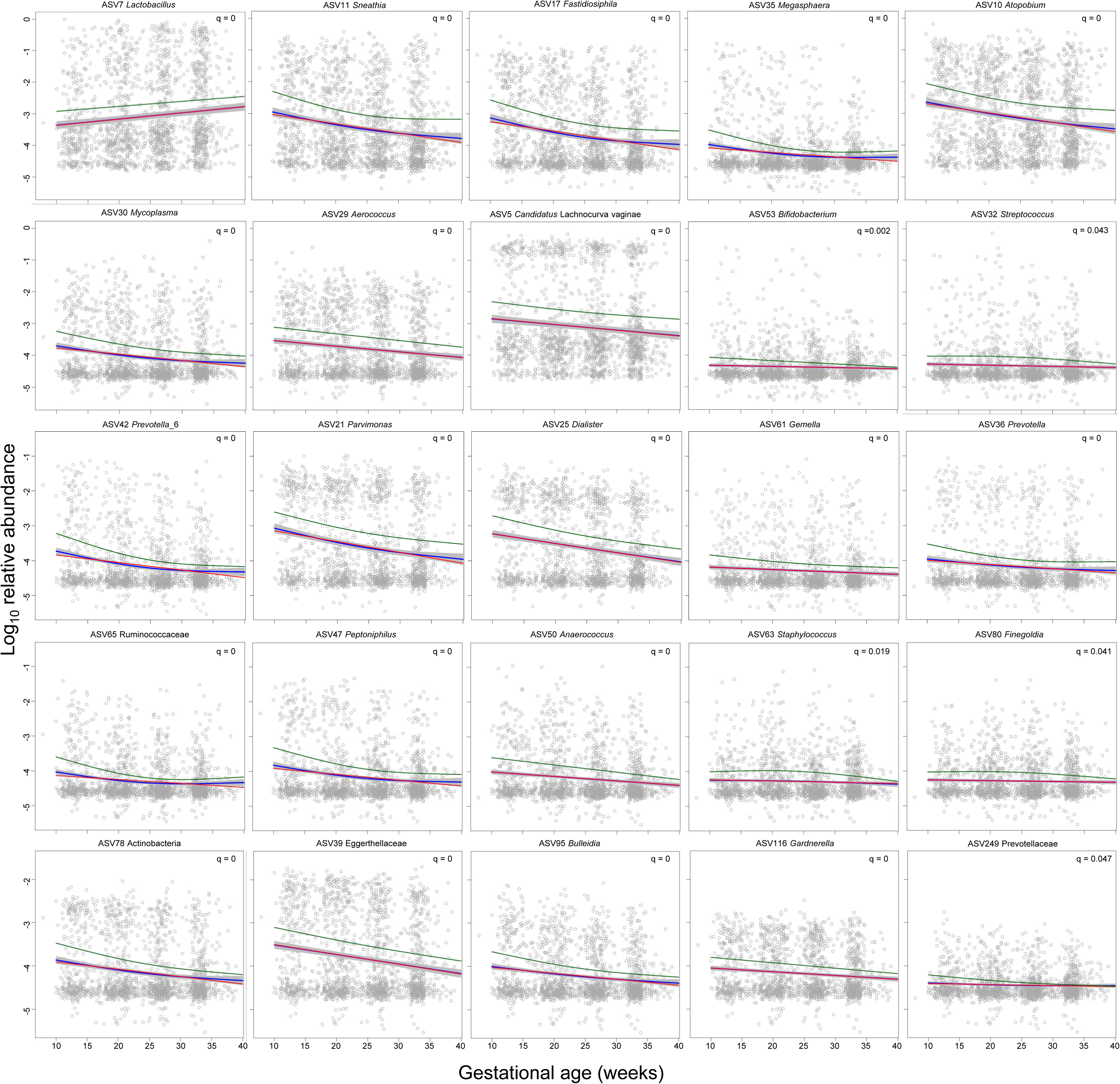
Changes in the relative abundance of amplicon sequence variants (ASVs) in vaginal 16S rRNA gene profiles across gestational age in women who ultimately delivered at term. Only the first ASV for each microbial taxon with a significant corrected p-value (q<0.05) presented in Supplemental Table 1 is shown. Panels with positive correlations are ordered before those with negative correlations. Each dot within an individual panel corresponds to one sample. The red lines represent linear fits through relative abundance data using linear mixed-effects models. The blue lines and grey bands represent the model fits and 95% confidence intervals derived from generalized additive models, respectively. The green lines represent the estimates from negative binomial mixed-effects generalized additive models.

To supplement the LME models at the ASV level, we further implemented Analysis of Composition of Microbiomes, or ANCOM (101), to identify ASVs changing in abundance throughout gestation. Gestational age was treated as a main fixed effect, patient identity as a random effect, and maternal age, parity, *Cannabis* use, ethnicity, and race were included as covariates. Seventy-five ASVs were positively or negatively associated with gestational age **(Supplemental Table 2, Figure 6)**. Fifty-six of these ASVs overlapped with those identified as being significantly associated with gestational age in the LME analysis **(Supplemental Figure 1)**. As in the LME analysis, many *Lactobacillus* ASVs were positively associated with gestational age while many bacteria typically associated with CST IV were negatively associated with gestational age. The direction of these associations is intriguing given prior reports that *Lactobacillus*-dominated vaginal CSTs are linked with positive reproductive outcomes (23, 26, 28) while, conversely, CST IV and CST IV-typical bacteria have been associated with an increased risk of sPTB (25, 28, 38). Notably, however, the ANCOM analyses additionally indicated that multiple ASVs classified as Ca. Lachnocurva vaginae also increased in abundance with advancing gestation. Ca. Lachnocurva vaginae, previously referred to as *Shuttleworthia* spp. (12), is an established resident bacterium of the vaginal ecosystem (102, 103), and it is typically a component of CST IV. Its increase in abundance throughout pregnancy is a novel finding, the potential clinical significance of which warrants further investigation.

**Figure 6.**
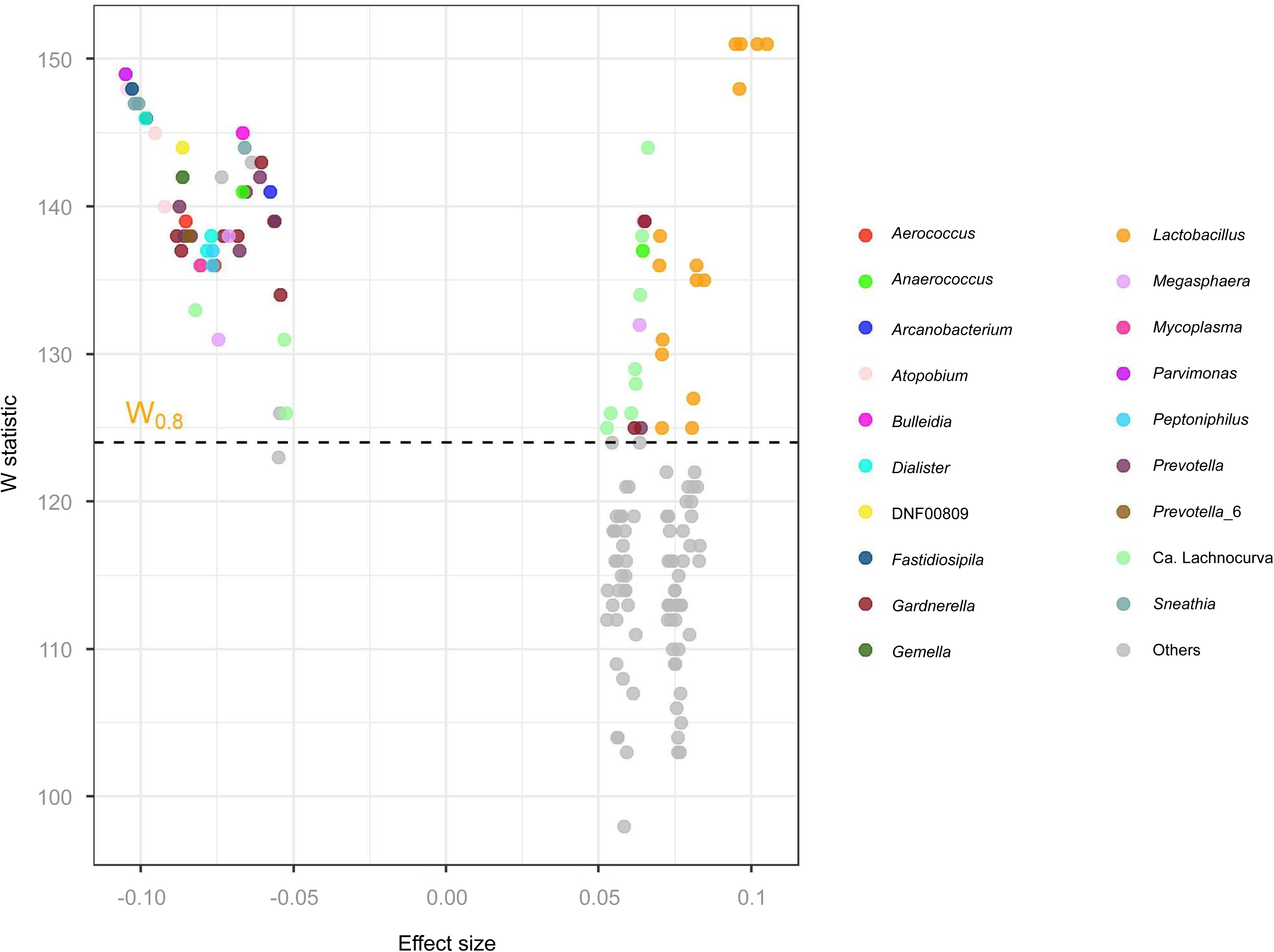
Amplicon sequence variants (ASVs) classified at the genus level which were identified as being less or more abundant in the vaginal microbiota with advancing gestational age. As gestation advances, *Lactobacillus*, and to a lesser extent Ca. Lachnocurva, ASVs become more abundant and many members of community state type (CST) IV become less abundant.

Overall, the results were largely congruent between the linear mixed-effects models and ANCOM analyses, although there were some notable differences (e.g. Ca. Lachnocurva vaginae). The fundamental difference between LME and ANCOM is that ANCOM is inferring about abundances whereas LME is inferring about relative abundances. Specifically, LME evaluates whether the abundance of a particular taxon, in a unit volume of an ecosystem, relative to all other taxa, has changed between two ecosystems. On the other hand, ANCOM evaluates whether the abundance of a particular taxon, in a unit volume of an ecosystem, has changed between two ecosystems. This explains the differences in the results obtained using the two approaches. Regardless, LME and ANCOM identified ecologically plausible variation in microbiota membership across gestational age, and a large proportion of the bacterial ASVs changing in composition and abundance across pregnancy were discovered using both approaches.

### Effect of maternal parity

In addition to the changes in alpha and beta diversity observed with increasing gestational age, there was also a significant effect of parity. Specifically, parity was positively correlated with alpha diversity (**Table 2**). One potential explanation for the association between parity and increased alpha diversity is that there is a marked increase in the alpha diversity of the vaginal microbiota after live birth (85, 104), and this phenomenon may be cumulative across multiple pregnancies, mirroring maternal-fetal immunological memory (105–108). Notably, this phenomenon cannot be explained simply by advancing maternal age, since this covariate was not positively correlated with alpha diversity of the vaginal microbiota (**Table 2**). This finding indicates that, while there is a consistent reduction in the richness and evenness of the vaginal microbiota throughout pregnancy, at least among women who have a diverse microbiota at pregnancy onset, this effect may be mitigated by parity.

With respect to beta diversity, only modest percentages of the variance in the composition and structure of the vaginal microbiota were explained by parity (0.3%-1.9%). Nevertheless, differences in CST membership based on parity, while adjusting for gestational age, were found (**Table 3**), with parity (OR=1.46 for each additional previous delivery) being associated with a decrease in CST III. At an ASV-level, higher maternal parity was significantly associated with an increase in 58 ASVs classified as typical vaginal CST IV bacteria (e.g., *Gardnerella*, *Megasphaera*, *Prevotella*, and *Sneathia*), while lower maternal parity was exclusively correlated with 7 *Lactobacillus* ASVs, 6 of which were classified as *L. crispatus* (q<0.1) (**Supplemental Table 1**). This finding is consistent with a recent report of increased vaginal microbiota diversity with higher parity during subsequent gestations (109). The ecological and clinical implications of the correlation between increased parity and vaginal microbiota diversity warrant further investigation.

**Table 3.**
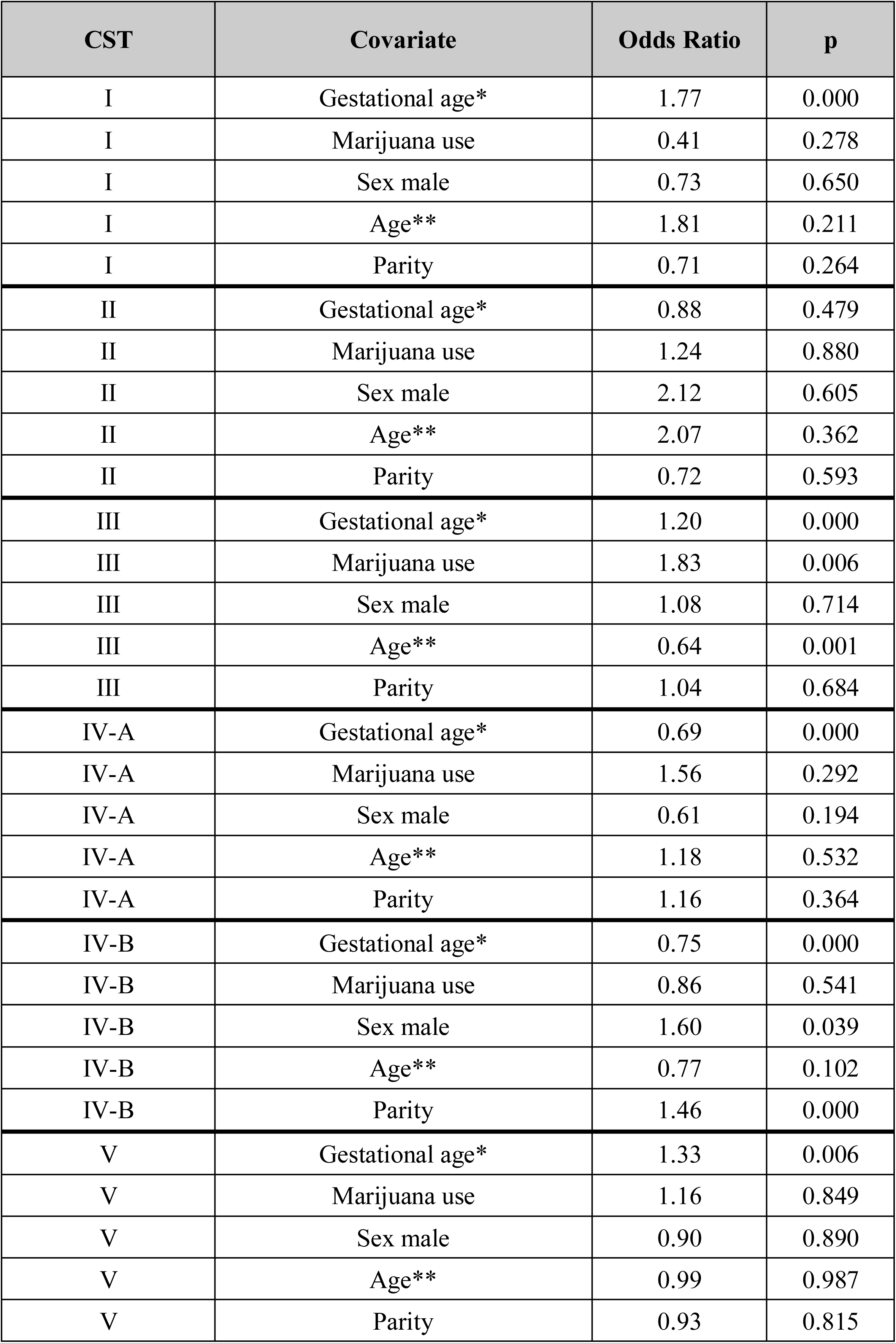
Factors associated with variation in vaginal community state type (CST) membership among women who delivered at term. *Gestational age was scaled by 4, therefore the Odds Ratio corresponds to a four week interval **Maternal age was scaled by 5, therefore the Odds Ratio corresponds to 5 year increase in maternal age

### Effect of maternal age

Only modest percentages of the variance in the composition and structure of the vaginal microbiota were explained by maternal age (0.2%-1.4%). Nevertheless, differences in CST membership based on maternal age, while adjusting for gestational age, were found (**Table 3**), with higher maternal age (OR=0.64 for each additional 5 years) being associated with a decrease in CST III. Similarly, at the ASV level, there were significant negative correlations between maternal age and 18 ASVs, 16 of which were classified as *L. iners*, while only 4 ASVs, classified as *L. crispatus*, were positively correlated with maternal age (**Supplemental Table 1**). While these correlations contrast with a previous report (47), the differences in ethnic makeup and sample size between the two cohorts could account for this discrepancy.

### Effect of obesity

There was a significant effect of obesity, defined as having a BMI greater than 28 kg/m^2^, on alpha diversity of the vaginal microbiota (**Table 2**). Specifically, there was a positive correlation between obesity and richness of the vaginal microbiota across gestation. This is in contrast with the intestinal microbiota, for which there tends to be a negative correlation between obesity and richness across gestation (110). Similar patterns are evident outside pregnancy as well. Obesity is associated with high alpha diversity of the vaginal microbiota (111) and, in general, low alpha diversity of the gut microbiota (112–115). Thus, for both of these body sites, obesity is associated with levels of microbiota alpha diversity that are widely viewed as non-optimal. As obesity is characterized by a low-grade systemic inflammatory response (116–119), these data highlight potential dynamic interactions between systemic inflammation and microbiota alpha diversity throughout the human body that can influence health and disease, including pregnancy outcomes (120, 121).

### Effect of Cannabis use

Only modest percentages of the variance in the composition and structure of the vaginal microbiota were explained by self-reported *Cannabis* use (0.3%-1.8%). Nevertheless, *Cannabis* use was associated with an increase in ASVs classified as *L*. *iners* (16 ASVs) and a decrease in those classified as *L*. *crispatus* (6 ASVs) (**Supplemental Table 1**). Given that *L. crispatus* has been associated with female reproductive health and positive pregnancy outcomes (4, 13, 15, 16, 19, 20, 25, 27, 28, 36, 40, 44, 89, 90, 92, 97, 122–127), these findings, among other potential general concerns (128, 129), caution against *Cannabis* use during pregnancy.

### Bacterial taxa are highly correlated with one another during normal pregnancy

**Supplemental Table 3** shows some of the strongest associations (LME adjusted q<0.05 and absolute spearman correlation coefficient >0.5) between pairs of bacterial taxa during pregnancy. In this analysis, each genus-level taxon (or family, if genus-level designation was not available) was represented by one ASV retained based on the strongest association with gestational age. A subset of these significant correlations (involving the most relatively abundant ASVs) is shown in **Figure 7**. *Atopobium* (ASV10) and *Gardnerella* (ASV3) (r=0.74), Eggerthellaceae (ASV39) and *Parvimonas* (ASV21) (r=0.83), *Dialister* (ASV25) and Eggerthellaceae (ASV39) (r=0.83), and *Sneathia* (ASV11) and *Parvimonas* (ASV 21) (r=0.74) were among the most highly correlated pairs of bacterial taxa in pregnancy (**Supplemental Table 3**). These data suggest potential synergistic relationships among these typical members of CST IV in pregnancy and potentially beyond.

**Figure 7.**
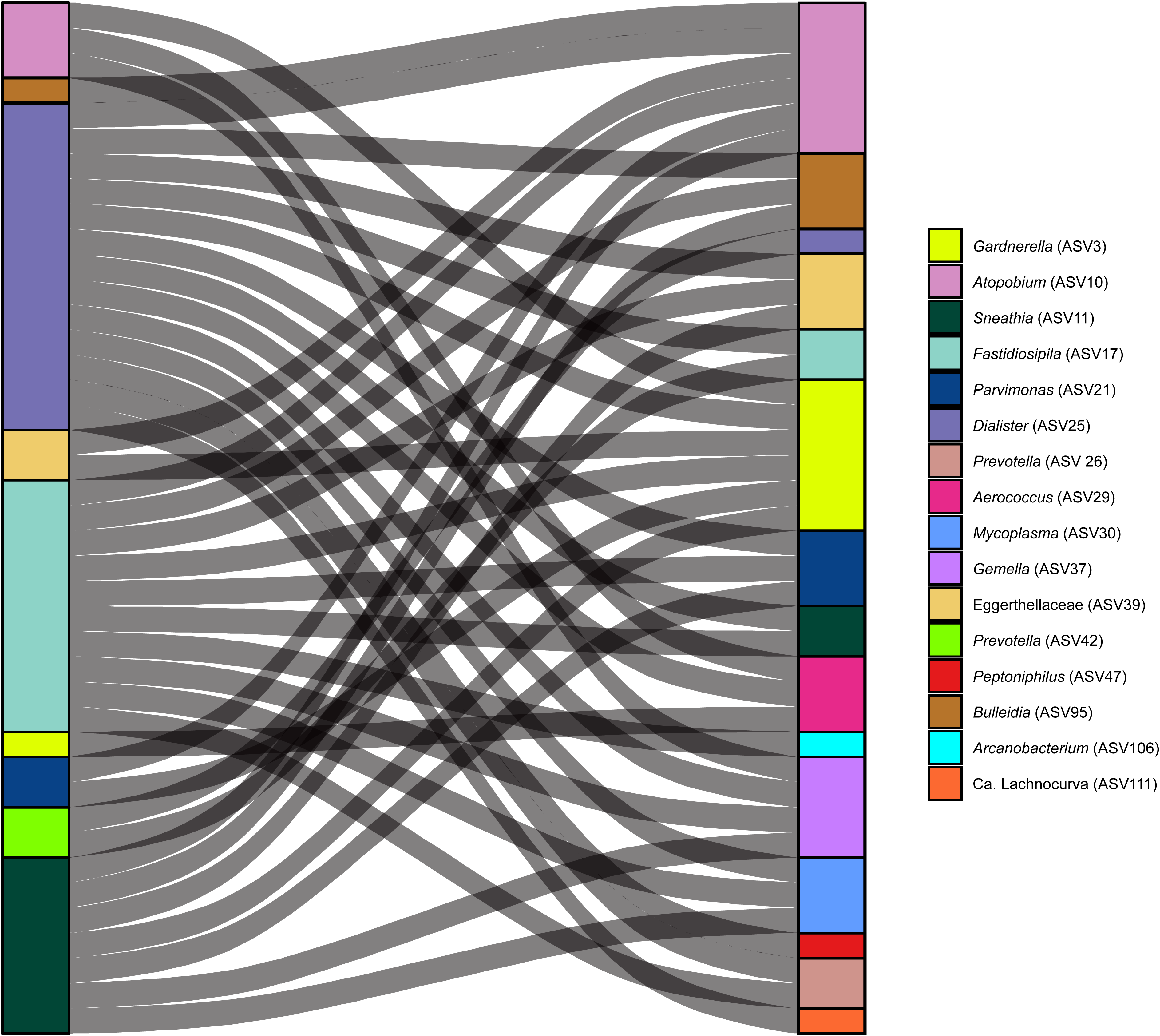
Positive correlations of relative abundances of vaginal microbial taxa. Alluvial plot shows pairs of vaginal bacterial taxa with highly correlated relative abundances throughout gestation. Relative abundances of amplicon sequence variants (ASVs) were compared using linear mixed-effects models.

### Network analysis reveals further changes in microbiota structure throughout pregnancy

The results from LME modeling were followed up with network analyses throughout gestation. Network analyses of the 25 most relatively abundant ASVs revealed that *Lactobacillus* ASVs were consistently network hubs, defined as ASVs closest to the center of the network, across term gestation **(Figure 8A-D)**. It is worth mentioning that there was limited resolution to differentiate *Lactobacillus* spp., given that the V4 hypervariable region of the 16S rRNA gene was targeted for sequencing (130). Nevertheless, *Lactobacillus* ASVs were clearly split between a primary group (ASVs 2, 7, 12, 13, 15, and 20) that included a mixture of *L. crispatus, L. jensenii,* and *L. gasseri*, and a secondary group (ASVs 1 and 6) comprised exclusively of *L. iners* **(Figure 8A-D)**. These positive associations were interesting, given the exclusionary nature of the CST-defining *Lactobacillus* spp. in the former group. In addition, this group maintained strong negative associations with *Gardnerella* (ASVs 3, 8, and 9), *Atopobium* (ASV 10), and *Megasphaera* (ASV 4) throughout gestation. By contrast, *L. iners* ASVs had very few associations with other ASVs, either positive or negative **(Figure 8A-D)**.

**Figure 8.**
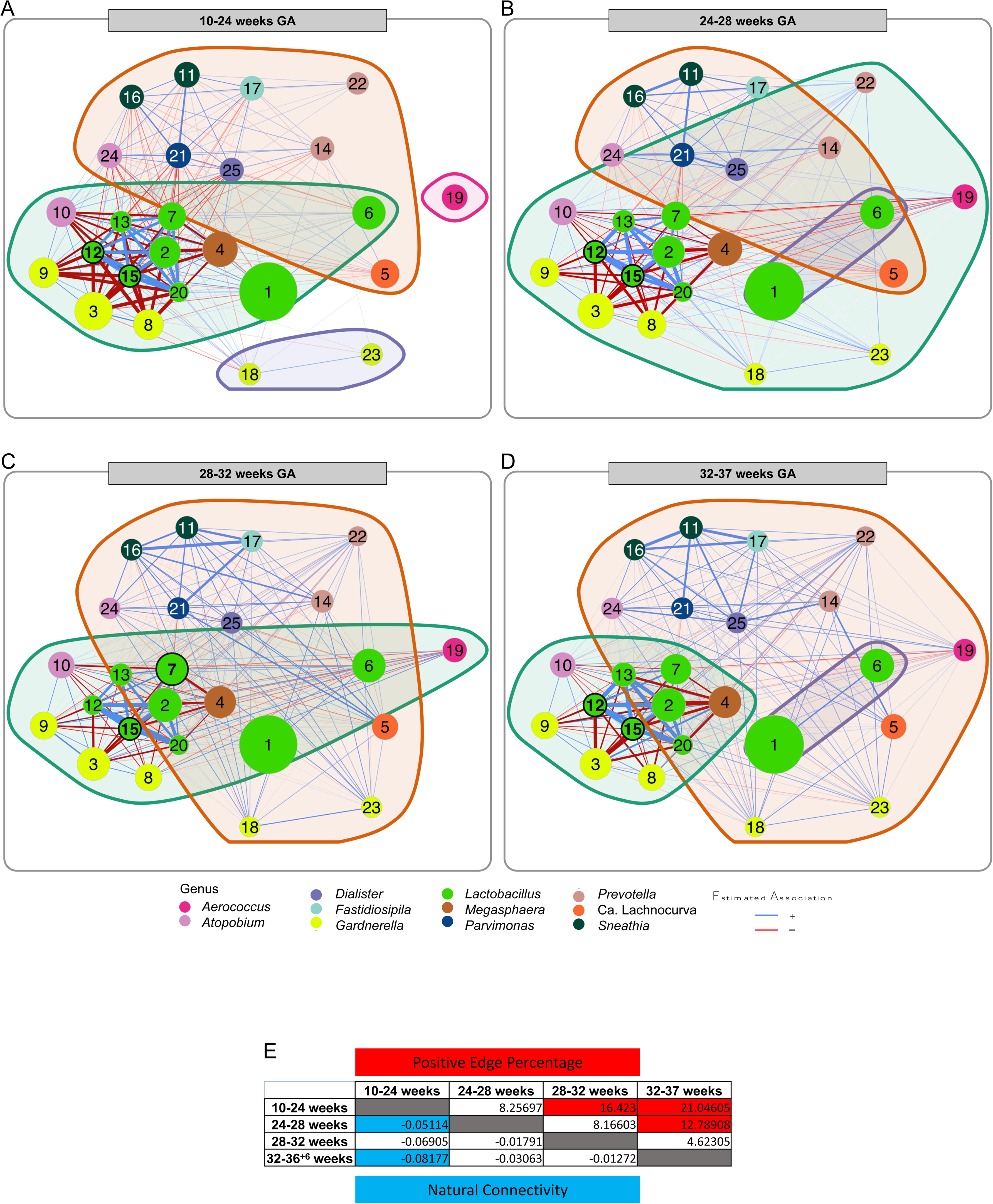
Network analysis illustrating changes in associations between amplicon sequence variants (ASVs) throughout pregnancy. Networks at (A) 10-24 weeks, (B) 24-28 weeks, (C) 28-32 weeks, and (D) 32-37 weeks of gestation were generated using the NetCoMi package (149). Nodes, which represented individual amplicon sequence variants (ASVs) were color coded according to their respective genus-level classification. Edges were weighted by strength using fitness and color coded by interaction type with positive (blue) and negative (red) interactions. Nodes that represent hubs, defined as an ASV with an eigenvector above the 95% quantile of the empirical distribution, are outlined in black and are in bold font. Clusters are represented by background coloration and darker borders. (E) A matrix of comparative network statistics with positive edge percentage above the diagonal and natural connectivity below the diagonal. Cells shaded red and cells shaded blue represent statistically significant differences in the respective time periods for positive edge percentage and natural connectivity, respectively.

This dichotomy may be due to the species-specific ability of *Lactobacillus* to produce lactic acid (both L- and D- isomers) and - to a lesser extent - hydrogen peroxide, each of which can create hostile conditions for other bacteria (131, 132), While *L. iners* can produce L-lactic acid, it lacks key genes to produce D-lactic acid (133). Conversely, *Lactobacillus crispatus* produces both isomers (127). Given that these two isomers of lactic acid differentially affect the biochemistry of vaginal fluid (127), they may also differentially influence the composition of the broader microbiota. Furthermore, unlike *L. crispatus*, *L. iners* lacks the ability to produce hydrogen peroxide (20, 134). Hydrogen peroxide is an established antimicrobial compound, yet it may only play a minor role within the vaginal ecosystem, as its production may be limited in this typically hypoxic environment (135). Regardless, the inability to produce both inhibitory metabolites may explain the lack of strong negative associations, and therefore, the permissive nature of *L. iners*, towards other ASVs when it predominates in the vagina (20).

We further analyzed the networks by defining clusters, formed by optimal grouping of ASVs based on strengths of association, which revealed differences in the degree of association between *Lactobacillus* spp. and CST IV-typical bacteria **(Figure 8A-D)**. In particular, CST IV bacteria were split between two groups. The first, which contained *Gardnerella* (ASVs 3, 8, 9), *Atopobium* (ASV 10), and *Megasphaera*, exhibited strong negative associations with *Lactobacillus* ASVs **(Figure 8A-D)**. The second, which contained *Sneathia*, *Dialister*, *Fastidiosipila*, *Shuttleworthia*, *Parvimonas,* and *Atopobium* (ASV 24), had only weak negative associations with *Lactobacillus* ASVs **(Figure 8A-D)**. Interestingly, bacteria within the latter group formed increasingly positive associations amongst themselves as gestation progressed **(Figure 8A-D)**. These contrasting patterns among non-*Lactobacillus* ASVs are in concordance with a prior report (104) that identified strong exclusionary associations between *L. crispatus* and *G. vaginalis* but only moderate negative associations between *L. crispatus* and other CST IV bacteria.

These clustering profiles mirror previously proposed splits within CST IV, with the green cluster comprised of *Gardnerella, Atopobium,* and *Megasphaera* representing CST IV-B and the orange cluster, formed by diverse bacteria (*Atopobium*, *Dialister*, Ca. Lachnocurva, *Parvimonas*, *Prevotella*, *Sneathia*) representing CST IV-C **(Figure 8A-D)**. While CST IV-B is defined by *Gardnerella* predominance, CST IV-C lacks a predominance of both *Lactobacillus* and *Gardnerella.* Instead, CST IV-C is formed by a multitude of diverse bacteria (11). The increase in positive associations amongst these particular CST IV-C members suggests that CST IV-C bacteria can co-exist, leading to more species-rich and diverse vaginal microbiotas than the more exclusionary CSTs. These findings are echoed in **Supplemental Table 3**, which reveals that the strongest associations among ASVs, as determined by LME modeling, exist among CST IV bacteria. For example, *Sneathia* ASV 11 and *Parvimonas* ASV 21 were highly correlated (r=0.74) and were consistently positively associated throughout gestation in the network analyses **(Figure 8A-D)**.

Intriguingly, G*ardnerella* ASVs 18 and 23 exhibited a distinct *Gardnerella-*correlative phenotype from the other *Gardnerella* ASVs (3, 8, and 9), which were in a separate cluster throughout most of gestation **(Figure 8 A,C-D)**. Instead of exhibiting the strong *Lactobacillus*-negative associations of *Gardnerella* ASVs 3, 8, and 9, G*ardnerella* ASVs 18 and 23 were often positively correlated with other ASVs throughout gestation, including those classified as *Lactobacillus* **(Figure 8B-D)**. Notably, *Gardnerella* ASV G2, which was an ASV associated with sPTB in a prior study (25), shared 100% identity with *G. vaginalis* ASV 9 and, in our study, it displayed strong negative associations with the *L. crispatus* cluster **(Figure 8A-D)**. ASVs 18 and 23 did not match any current *Gardnerella* type strain with 100% identity using BLAST **(Supplemental Table 4)**; they may represent unique *Gardnerella* strains. This is important because *Gardnerella* is associated with bacterial vaginosis (136) and sPTB (137), yet, seemingly in a strain-dependent manner (25). Therefore, these network analyses highlight the need for strain-level resolution of the vaginal microbiota to fully understand its complex dynamics and ecology in health and disease.

Lastly, each of the four networks from different time points in gestation were compared with each other to identify global and ASV-specific network changes. Globally, natural connectivity (i.e., robustness of the network) significantly decreased and positive edge percentage (i.e., proportion of positive associations) significantly increased **(Figure 8E)** as pregnancy progressed, both with strong linear trends across gestation (R^2= 0.895 and 0.985, respectively). The increase in positive edge percentage was a combination of a decrease in negative *Lactobacillus*-CST IV-B associations, and an increase in the strength of positive associations among CST IV-C-associated ASVs. These gestational changes observed in positive edge percentage were associated with a significant decrease in closeness (i.e., the sum of the shortest paths between a node and all other nodes) for *Gardnerella* ASVs 3, 8, and 9, since many of their strong negative *Lactobacillus* spp. associations were lost or diminished as pregnancy progressed (**Figure 8B-D)**. Conversely, positive associations for CST IV-associated ASVs increased in general, likely resulting from an increase in available niches within the vaginal ecosystem as term approaches. Such changes may be due to the shift towards *Lactobacillus-*dominated CSTs observed in **Figures 4B and 4C**.

Collectively, these network analyses demonstrate the complex interactions between members of the vaginal microbiota. Not only were we able to confirm classical associations of *Lactobacillus* species with members of other genera, but through longitudinal collection of vaginal samples, we identified shifts in network connectivity as gestation progressed. Furthermore, different ASVs attributed to the same genus (e.g., *Gardnerella*) demonstrated distinct ecological dynamics, suggesting that strain-level variation is indeed driving community phenotypes commonly denoted as CSTs. Therefore, this study highlights the need for strain-level investigations utilizing metagenomic data to further characterize these shifts in vaginal microbiota ecology throughout gestation and determine their underlying causes and consequences.

### Conclusions and Future Directions

The composition of the vaginal microbiota is broadly consistent across populations of reproductive age women worldwide (1–5), yet, the relative abundances of CSTs can vary within populations, including by ethnicity (2–5), and within individuals over time. It is increasingly hypothesized that variation in the vaginal microbiota is contributing to obstetrical complications, especially sPTB (22, 23, 25, 27, 28). Here, we have provided a longitudinal study of a primarily African-American population with a large sample size, and extensive demographic and clinical data, which allowed for the simultaneous evaluation of a broad range of maternal characteristics on the vaginal microbiota. Indeed, this study represents the largest and most comprehensive longitudinal survey of the vaginal microbiota throughout gestation resulting in a term delivery and thereby provides foundational understanding. The focus on African-American women is a clear strength of the study because they constitute a high-risk population that experiences a relatively high rate of pregnancy complications (65–75).

In the current study, we report that the principal factors influencing the composition and structure of the vaginal microbiota in pregnancy are individual patient identity and gestational age at sampling. The pronounced effect of individual identity highlights the need for longitudinal studies to account for inter- and intra-individual variability when evaluating the strengths of potential relationships between the composition of the vaginal microbiota and obstetrical complications. Furthermore, the richness and evenness of the vaginal microbiota decreased throughout pregnancy, with the microbiota becoming increasingly predominated by *Lactobacillus* species with advancing gestation. Such typical changes can be partially mitigated by maternal parity and obesity. Importantly, *Lactobacillus* species, especially *L. crispatus*, are generally perceived to promote the vaginal and reproductive health of women (4, 13, 15, 16, 19, 20, 25, 27, 28, 36, 40, 44, 89, 90, 92, 97, 122–127); therefore, any factors that could potentially reduce the likelihood of the transition of the vaginal microbiota to a *Lactobacillus*-dominant community during pregnancy need to be identified. Lastly, network analyses revealed dynamic interactions among individual bacterial strains within the vaginal microbiota during pregnancy and the number and structure of these interactions change with advancing gestation. A critical consideration moving forward will be to assess whether patterns in these strain-level interactions within the vaginal microbiota differ between women delivering at term or those who ultimately experience sPTB. Ideally, this should be done using metagenomics (138), in addition to 16S rRNA gene sequencing, so that strain-level designation of bacterial taxa can be more readily achieved and information on the functional and virulence potential of various strains within individual microbiotas can be gleaned (28, 104). Furthermore, a key element missing from most studies is the characterization of host-immune microbiome interactions, which can be readily assessed by evaluating the immunoproteome (cytokines, chemokines, defensins, etc.) within the vaginal ecosystem. The local immune responses of women to their vaginal microbiota may be as or more variable than the compositions of their microbiota themselves. Therefore, elucidating the dynamics of host immune-microbiome interactions and their potential influence on obstetrical outcomes, especially sPTB, is a critical future direction (27, 28, 139, 140).

## MATERIALS AND METHODS

### Vaginal fluid specimens

Vaginal fluid samples were obtained at the Perinatology Research Branch, an intramural program of the *Eunice Kennedy Shriver* National Institute of Child Health and Human Development, National Institutes of Health, U.S. Department of Health and Human Services, Wayne State University (Detroit, MI), and the Detroit Medical Center (Detroit, MI). The collection and use of human materials for research purposes were approved by the Institutional Review Boards of the National Institute of Child Health and Human Development and Wayne State University (#110605MP2F(RCR)). All participating women provided written informed consent prior to sample collection.

### Study design

This was a retrospective longitudinal cohort study to characterize variation in the vaginal microbiota across gestation in pregnancies ending in normal term delivery. A normal pregnancy was defined as a woman with no obstetrical, medical or surgical complications, who agreed to participate in this study, provided written signed informed consent, and delivered at term (38 to 42 weeks) without complications. Three or four samples of vaginal fluid were collected longitudinally across pregnancy from each woman under direct visualization from the posterior vaginal fornix using a Dacron swab (Medical Packaging Corp., Camarillo, CA). Vaginal swabs were stored at −80°C until time of DNA extraction.

### DNA extraction from vaginal swabs

Genomic DNA was extracted from vaginal swabs (N=1,862) alongside non-template negative controls addressing any potential background DNA contamination (N=73). All vaginal swabs were randomized across extraction runs. Extractions were conducted using a Qiagen MagAttract PowerMicrobiome DNA/RNA EP extraction kit (Qiagen, Germantown, MD), with minor modifications to the manufacturer’s protocols. Briefly, swabs were transferred to clean, labeled Corning cryovials (Corning, Corning, NY) and immersed in 750 µL solution MBL pre-heated to 60°C. Swabs were then vortexed for 10 min. A provided empty PowerBead plate was then centrifuged for 1 min at 4,400 x g, and vaginal swab lysates were added to corresponding wells of the PowerBead plate. Plates containing lysates were centrifuged for 1 min at 4,400 x g. The plates were then loaded onto a TissueLyser II plate shaker (Qiagen, Germantown, MD), firmly secured, and shaken at 17 Hz for 20 min. Plates were then removed from the shaker and immediately centrifuged at 4,400 x g for 6 min. The supernatant was then carefully transferred 185 µL at a time to a provided collection plate. Following transfer, 150 µL of solution IRS was added to each well and plates were incubated at 4°C for 10 min. Plates were centrifuged for 15 min at 4,400 x g and supernatant was transferred to a new collection plate. The plate was centrifuged for 2 min at 4,400 x g, and 850 µL of the supernatants were transferred to a clean collection plate. The collection plate was loaded onto the epMotion 5075 liquid handler (Eppendorf, Enfield, CT, USA) for further processing following the default onboard protocols. The above procedure yielded between 0.13 and 550 ng/µL purified DNA from the vaginal swabs as measured by a Qubit 3.0 fluorimeter and Qubit dsDNA assay kit (Life Technologies, Carlsbad, CA) following the manufacturer’s protocol. The purified DNA was transferred to the provided 96-well microplates and stored at -20°C.

### 16S rRNA gene amplicon sequencing and bioinformatic processing

The V4 region of the 16S rRNA gene was amplified from vaginal swab DNA extracts and sequenced at Michigan State University’s Research Technology Support Facility (https://rtsf.natsci.msu.edu/) using the dual indexing sequencing strategy developed by Kozich et al. (141). The forward primer was 515F: 5’-GTGCCAGCMGCCGCGGTAA-3’ and the reverse primer was 806R: 5’-GGACTACHVGGGTWTCTAAT-3’. Each PCR reaction contained 0.5 µM of each primer, 1.0 µl template DNA, 7.5 μl of 2X DreamTaq™ Hot Start PCR Master Mix (Life Technologies, Carlsbad, CA), and nuclease-free water to produce a final volume of 15 µl. Reactions were performed using the following conditions: 95 °C for 3 minutes, followed by 30 cycles of 95 °C for 45 seconds, 50 °C for 60 seconds, and 72 °C for 90 seconds, with an additional elongation at 72 °C for 10 minutes.

16S rRNA gene amplicon sequences were clustered into amplicon sequence variants (ASVs) defined by 100% sequence similarity using DADA2 version 1.12 (142) in R version 3.6.1 (143) according to the online MiSeq protocol (https://benjjneb.github.io/dada2/tutorial.html) with minor modifications, as previously described (144). These modifications included allowing truncation lengths of 250 and 150 bases, and a maximum number of expected errors of 2 and 7 bases, for forward and reverse reads, respectively. Reads were truncated at the first instance of a quality score less than or equal to 2. Any reads containing ambiguous nucleotides were removed from the dataset. To increase power for detecting rare variants, sample inference allowed for pooling of samples. Additionally, samples in the resulting sequence table were pooled prior to removal of chimeric sequences. Sequences were then classified using the silva_nr_v132_train_set database with a minimum bootstrap value of 80%, and sequences that were derived from Archaea, chloroplast, or Eukaryota were removed. Per Holm et al. (12), ASVs classified as *Shuttleworthia* were manually reclassified as Ca. Lachnocurva vaginae.

The R package decontam version 1.6.0 (145) was used to identify ASVs that were likely potential background DNA contaminants based on their distribution among biological samples and negative controls using the “IsContaminant” method. An ASV was identified as a contaminant and subsequently removed from the dataset if it had a decontam P score ≤ 0.5, was present in at least 15% of negative controls with an overall average relative abundance of at least 1.0%, and had a greater average relative abundance in controls than biological samples. Based on these criteria, a total of four ASVs classified as *Escherichia*, *Pelomonas*, *Pseudomonas*, and Micrococcaceae were identified as contaminants.

For assigning community state types (CSTs) to the bacterial community profiles, ASVs were first taxonomically classified using the V4_trimmed_noEuks_nr_Complete.fa reference library supplied with the speciateIT classifier code (http://ravel-lab.org/speciateit/) and the classify.seqs command in mothur (146) with a bootstrap cutoff value of 80. Read counts for ASVs that were assigned to the same taxon were then combined, and CSTs were assigned using VALENCIA, a nearest centroid-based classifier (11).

### Statistical analysis

#### Analysis of vaginal microbiota composition and structure

To determine the percentage of variance explained (R^2^) in the composition (Jaccard index) or structure (Bray-Curtis index) of the vaginal microbiota, PERMANOVA analyses (147) were performed using the interaction terms between SubjectID and gestational age at sampling within the “adonis2” function in the R package vegan version 2.5-6 (148). The confidence intervals of R^2^ statistics were obtained by bootstrap sampling of patients and all their associated longitudinal measurements. Confidence intervals for R^2^ statistics for additional patient specific covariates (i.e., maternal age, parity, obesity, race, ethnicity, *cannabis* use) while accounting for gestational age were obtained in a separate analysis in which PERMANOVA analysis was performed on bootstrap samples of subjects. For each subject only one random longitudinal observation was selected, hence generating cross-sectional datasets in which observations were independent and hence PERMANOVA could be applied. Empirical 95% confidence intervals of R^2^ statistics were obtained using 1000 bootstrap iterations.

#### Changes in alpha diversity with gestational age and maternal characteristics

Relative abundance for each amplicon sequence variant (ASV) was determined as the ratio of the count of each ASV divided by the total number of ASVs in each sample. Starting with the relative abundance data, for each sample, we calculated the Shannon and Simpson diversity using the *diversity* function in the *vegan* package and the Chao1 diversity using the *chao1* function in the *fossil* package. Each measure of diversity was then correlated with the continuous variable gestational age at sampling using linear mixed-effects models implemented in the *lme4* package in R. In these models a random intercept and a random slope with gestational age was allowed for each subject to account for the repeated and potentially correlated observations from the same subject. Gestational age values were centered at 8 weeks and then scaled by 4 to facilitate interpretation of random intercepts and convergence of model fitting algorithms. The complexity of the gestational age dependence was assessed by comparing the model fit between a linear and quadratic trend using a likelihood ratio test for linear mixed-effects models. The same test was used to determine the need for subject specific gestational age slopes. To further inspect nonlinear trends in alpha diversity as a function of gestational age, Generalized Additive Models (GAM) for repeated observations were also fit based on spline transformation of gestational age. Such models were available from the *mgcv* package in R. The effects of maternal characteristics (maternal age, obesity, parity, race, ethnicity, smoking, and *cannabis* use) were assessed by including these as co-variates in linear mixed-effects models. A p-value <0.05 was used to infer significance in these analyses.

#### Changes in vaginal community state types (CSTs) with gestational age and maternal characteristics

The log-odds of membership in a given community state type (CST) were modeled using binomial linear mixed-effects models using the *glmer* function in R. Fixed effects in these models included gestational age at sampling (linear and quadratic terms, as needed) and maternal characteristics. Of note, due to the sparse responses in these models (membership to a given CST), it was not feasible to test whether there were subject-specific departures in the CST membership probability trends versus gestational age (random slopes for gestational age), yet subject specific shifts in membership probabilities were allowed via random intercepts in the mixed effects models.

#### Changes in the relative abundance of individual amplicon sequence variants (ASVs) with gestational age and maternal characteristics

The analysis of the relative abundance of each amplicon sequence variant (ASV) in association with gestational age at sampling was performed using linear mixed-effects models based on ASV count data while assuming a negative binomial distribution of the counts. Such models were implemented in the *glmmTMB* package in R and included an offset term of the total number of reads per sample, so that changes in relative abundance with gestational age were being estimated as opposed to differences in absolute counts. These models included gestational age and maternal characteristics as fixed effects and random intercept and random gestational age slope for each subject. All analyses involved control of the false discovery rate at a 10% level (q<0.1).

Additionally, we implemented Analysis of Composition of Microbiomes, or ANCOM (101), for further differential abundance analysis of ASVs. After adding a pseudo-count (1) to all observed abundances, ANCOM accounts for the compositionality issue of the microbiome data by performing the additive log ratio (ALR) transformation. For each taxon, ANCOM uses all other taxa, one at a time, as the reference in forming the ALR transformation. The transformed data were treated as the response of the LME model which includes gestational age as the main fixed effect, maternal age, parity, marijuana use, ethnicity, and race as covariates while allowing a random intercept and a random slope for each subject. For a given taxon, the output W statistic represents the number of ALR transformed models where the taxon is differentially abundant with regard to the main fixed effect, after adjusting for multiple testing correction for the number of ALR models corresponding to each taxon. The larger the value of W, the more likely the taxon is differentially abundant between compared sample groups.

#### Bacterial taxa with the most highly associated relative abundances during normal pregnancy

To assess the association between pairs of ASVs, we modeled their log-transformed relative abundance data using linear mixed-effects models. In these models one of the two ASVs was treated as a response variable while the other was treated as an explanatory variable. A random effect was allowed for each subject. Naïve Spearman correlation coefficients were also calculated for each pair. Significance of correlations was based on an adjusted p-value <0.05.

#### Associations of ASVs across term pregnancy through network analysis

The R packages NetCoMi (149)1.0.2, SpiecEasi 1.1.2, and seqtime 0.1.1 were used in R version 4.0.3 to create correlation networks between ASVs from vaginal samples across four time points in gestation resulting in term delivery. Only one sample per subject was included for each time point to control for subject ID. Networks were generated using Spearman’s correlation since the data were not normally distributed and nonparametric. Only the top 25 predominant ASVs in the entire dataset were considered for each network. A topological overlap matrix generated from the network adjacency matrix was utilized as a dissimilarity measure after transforming the data through multiplicative simple replacement and a centered log-ratio transformation to account for the zero-inflated data and to normalize the data, respectively. Community structure was determined by implementing the fast greedy modularity optimization algorithm (150). The layout of the network for the first time period was used as the layout for all subsequent time periods. Edges displayed in the network exceeded a threshold of 0.3 and edge thickness was tied to the strength of the correlation between two given nodes. Networks of the four time periods were compared using the NetCoMi package “netCompare” function with 1000 permutations.

## ACKNOWLEDGEMENTS

We thank the physicians and nurses from the Center for Advanced Obstetrical Care and Research and the Intrapartum Unit for their help in collecting human samples. The authors also thank the staff members of the PRB Clinical Laboratory for the processing of these samples. This research was supported, in part, by the Perinatology Research Branch, Division of Obstetrics and Maternal-Fetal Medicine, Division of Intramural Research, *Eunice Kennedy Shriver* National Institute of Child Health and Human Development, National Institutes of Health, U.S. Department of Health and Human Services (NICHD/NIH/DHHS) under Contract No. HHSN275201300006C. KRT, AT, and NG-L were further supported by the Wayne State University Perinatal Research Initiative in Maternal, Perinatal and Child Health. Dr. Romero has contributed to this work as part of his official duties as an employee of the United States Federal Government.

## SUPPLEMENTAL TABLE LEGENDS

**Supplemental Table 1. Multivariate analysis of relative abundances of amplicon sequence variants (ASVs) in vaginal 16S rRNA gene data as a function of gestational age, maternal age, parity, ethnicity, and marijuana use among women who ultimately delivered at term.**

The p-value and false discovery rate adjusted p-value (q-value) are provided for each covariate. The coefficients represent changes in log (base e) relative abundance with: (A) one additional month of gestational age, (B) 5 additional years of maternal age, (C) use of marijuana, (D) ethnicity Hispanic or Latino, (E) race other than African American, and (F) one additional previous delivery. ASVs classified at the genus level as *Lactobacillus* were secondarily classified at the species level, if possible, using the National Center for Biotechnology Information’s BLAST (Basic Local Alignment Search Tool). These ASVs are highlighted in blue.

**Supplemental Table 2. Comparison of analyses evaluating the relationship between gestational age and the vaginal microbiota using absolute abundance analysis (Analysis of Composition of Microbiomes; ANCOM) and relative abundance analysis (Linear mixed-effects models; LME).**

**Supplemental Table 3.** Vaginal bacterial taxa with highly correlated relative abundances based on longitudinal sampling in women who ultimately delivered at term.

Significance of correlations between log-transformed relative abundances was assessed using linear mixed-effects models. In these models one of the two amplicon sequence variants (ASVs) was treated as a response while the other as an explanatory variable. A random effect was allowed for each subject. Naïve spearman correlation coefficients were calculated for each pair of taxa. The p-value and false discovery rate-adjusted p-value (q-value) are provided.

**Supplemental Table 4. Top 25 amplicon sequence variants (ASVs) by relative abundance compared to NCBI BLAST bacterial type strains with percent identity.**

ASVs were compared to type strains in the NCBI BLAST database. Highest percentage identity type strains were listed if not 100%.

## SUPPLEMENTAL FIGURE LEGENDS

**Supplemental Figure 1. Venn diagram showing the relationship between absolute abundance analysis (Analysis of Composition of Microbiomes; ANCOM) and relative abundance analysis (Linear mixed-effects models; LME) in evaluating changes in the structure of the vaginal microbiota throughout gestation.** The two analyses identified some common significant bacterial taxa (i.e., amplicon sequence variants, or ASVs), yet the ANCOM approach tends to be more conservative than LME.

